# Cortical responses to looming sources are explained away by the auditory periphery

**DOI:** 10.1101/2023.11.30.569391

**Authors:** Sarah Benghanem, Rudradeep Guha, Estelle Pruvost-Robieux, Julie Lévi-Strauss, Coralie Joucla, Alain Cariou, Martine Gavaret, Jean-Julien Aucouturier

**Affiliations:** INSERM UMR 1266, IPNP (Institut de Psychiatrie et Neurosciences de Paris), Paris, France; Medical ICU, Cochin Hospital, AP-HP, Paris, France; University Paris Cité, Medical School, Paris, France; Université de Franche-Comté, SUPMICROTECH, CNRS, institut FEMTO-ST, F-25000 Besançon, France; Neurophysiology department, GHU Paris Psychiatrie et Neurosciences, Sainte Anne Hospital, F-75014 Paris; Science & Technologies of Music & Sound (STMS), IRCAM/CNRS/Sorbonne Université, F-75004 Paris, France

**Keywords:** event-related potentials, mismatch negativity, oddball paradigm, looming sounds, Cortical responses

## Abstract

A wealth of behavioral evidence indicate that sounds with increasing intensity (i.e. appear to be looming towards the listener) are processed with increased attentional and physiological resources compared to receding sounds. However, the neurophysiological mechanism responsible for such cognitive amplification remains elusive. Here, we show that the large differences seen between cortical responses to looming and receding sounds are in fact almost entirely explained away by nonlinear encoding at the level of the auditory periphery. We collected EEG mismatch negativity (MMN) data in response to deviant stimuli with both dynamic (looming and receding) and constant level (flat) differences to the standard in the same participants. We then combined a computational model of the auditory periphery with generative EEG methods (temporal response functions, TRFs) to model the single-participant MMN responses to flat deviants, and used them to predict the effect of the same mechanism on looming and receding stimuli. The flat model explained a remarkable 45% variance of the looming response, and 33% of the receding response. This provide striking evidence that MMN responses to looming and receding sounds result from the same cortical mechanism that generate MMN to constant-level deviants: all such differences are the sole consequence of their particular physical morphology getting amplified and integrated by peripheral auditory mechanisms. Thus, not all effects seen cortically proceed from top-down modulations by high-level decision variables, but can rather be performed early and efficiently by feed-forward peripheral mechanisms that evolved precisely to sparing subsequent networks with the necessity to implement such mechanisms.

## Introduction

The human auditory system has evolved to respond efficiently to fast and unpredictable changes in the acoustic environment that could be relevant for survival. One of the most salient examples of such prioritized auditory processing is the perceptual bias towards *looming* vs *receding* sound sources, which are typically simulated in the lab using simple increasing or decreasing changes of intensity sound level (Kolarik et al 2016(1)). The saliency of *looming* source produced by increasing intensity sound levels is a hallmark of human psychoacoustics: participants consistently overestimate the loudness (Neuhoff, 1998; Ponsot et al, 2015(2,3)) and speed (Rosenblum, Carello and Pastore, 1987, Schiff & Oldak, 1990(4,5)) of looming compared to receding sounds. Physiologically, looming sounds also elicit stronger orienting response measured by skin conductance and heart rate changes (Bach et al 2008; Bach et al, 2009, Tajadura-jimenez et al 2010(6–8)), and facilitate the processing of associated visual stimuli (Romei et al 2009, leo et al 2011(9,10)). Finally, brain imaging studies have shown that *looming* and *receding* sounds activate brain areas related to spatial auditory processing, which include the right temporal plane and the right superior temporal sulcus (Seifritz et al. 2002, Alho et al, 2014(11,12)), and that *looming* sounds activate a wider network of regions subserving auditory spatial perception and attention compared to receding sounds, including the right amygdala and left temporal areas (Seifritz et al. 2002, Bach et al., 2008(6,11)). In sum, a wealth of behavioral and brain-imaging evidence converges to indicate that sound with increasing intensity function is an elementary warning cue, able to elicit adaptive responses by recruiting additional attentional and physiological resources.

Despite all this, event-related potential (ERP) evidence for the prioritized or amplified processing of *looming vs receding* sounds have remained remarkably contrasted. When comparing changing-level sounds with more frequent constant-level standards, some studies have documented earlier and higher mismatch negativity (MMN) for looming than for receding sounds, which is coherent with the general pattern of “cognitive amplification” of looming sounds (Shestopalova et al 2018(13)); but others have found that MMN amplitude increase with the magnitude of intensity change irrespective of direction (Rinne et al, 2006(14)); and several studies have also reported no differences in MMN latencies or amplitudes for either *looming* or *receding* sounds (Altmann et al, 2013; Näätänen, 1992(15,16)).

On closer inspection, these different results were obtained with experimental stimuli which, despite sharing a general pattern of increasing or decreasing amplitude, actually display a wide diversity of temporal characteristics (stepwise or gradual changes, linear or exponential profiles, duration and onset of level ramp). Psychophysical research has highlighted that loudness integration in changing-level sounds is not identically distributed in time, and that perceptive weights are biased towards the beginning and end of the sound (Ponsot et al. 2013(17)). If ERPs reflect such integration, then the latency and amplitude of the MMN responses may depend heavily on the morphology of the deviant. In addition, electrophysiological research shows that, even in mice, populations of neurons in the auditory cortices respond to rising or decreasing intensity-ramps asymmetrically, as the direct result of neuronal adaptation and non-linearities in their temporal integration (Deneux et al. 2016 (18)). For all these reasons, very large differences between *looming vs receding* MMN responses do not necessarily indicate, as often implied in the literature, a top-down cognitive amplification or prioritization of one type of sound over the other, but could be the result of the bottom-up integration of complex temporal profiles in stimuli and temporal non-linearities in their subsequent processing, i.e. of the same generic sensory mechanisms that would generate MMN to e.g. unremarkable constant-level deviants.

To clarify whether and how ERP responses to *looming vs receding* sounds coincide with behavioral and fMRI evidence of their cognitive amplification, we need a way to model sensory contributions to MMN responses and to time-varying stimuli, and explore how much these generic mechanisms explain responses to specifically *looming* and *receding* sounds. To do this, we collected scalp EEG MMN data in response to deviant stimuli that presented potentially both dynamic (*looming* and *receding*) and constant level (flat) differences to the standard in the same participants. We then combined a model of the auditory periphery with generative EEG methods (temporal response functions, TRFs) to model the single-participant MMN responses to flat-intensity deviants, and used them to predict the effect of the same mechanism on time-varying looming and receding stimuli. By comparing actual vs predicted responses, we could investigate whether and how the EEG responses to looming sounds is specific to their arousing or salient nature or, on the contrary, explained away by generic auditory mechanisms.

## Results

We recorded 64-channel EEG from a sample of N=18 participants (9 females, median age = 25 years old) while they were presented a 30-minute sequence of frequent pure tones (1000Hz, 300ms) with constant-intensity combined with rare deviants that were both longer (600ms) and had either constant (flat) or dynamically changing intensity (*looming* and *receding*; see Methods & Figure 1.A).

**Figure 1:**
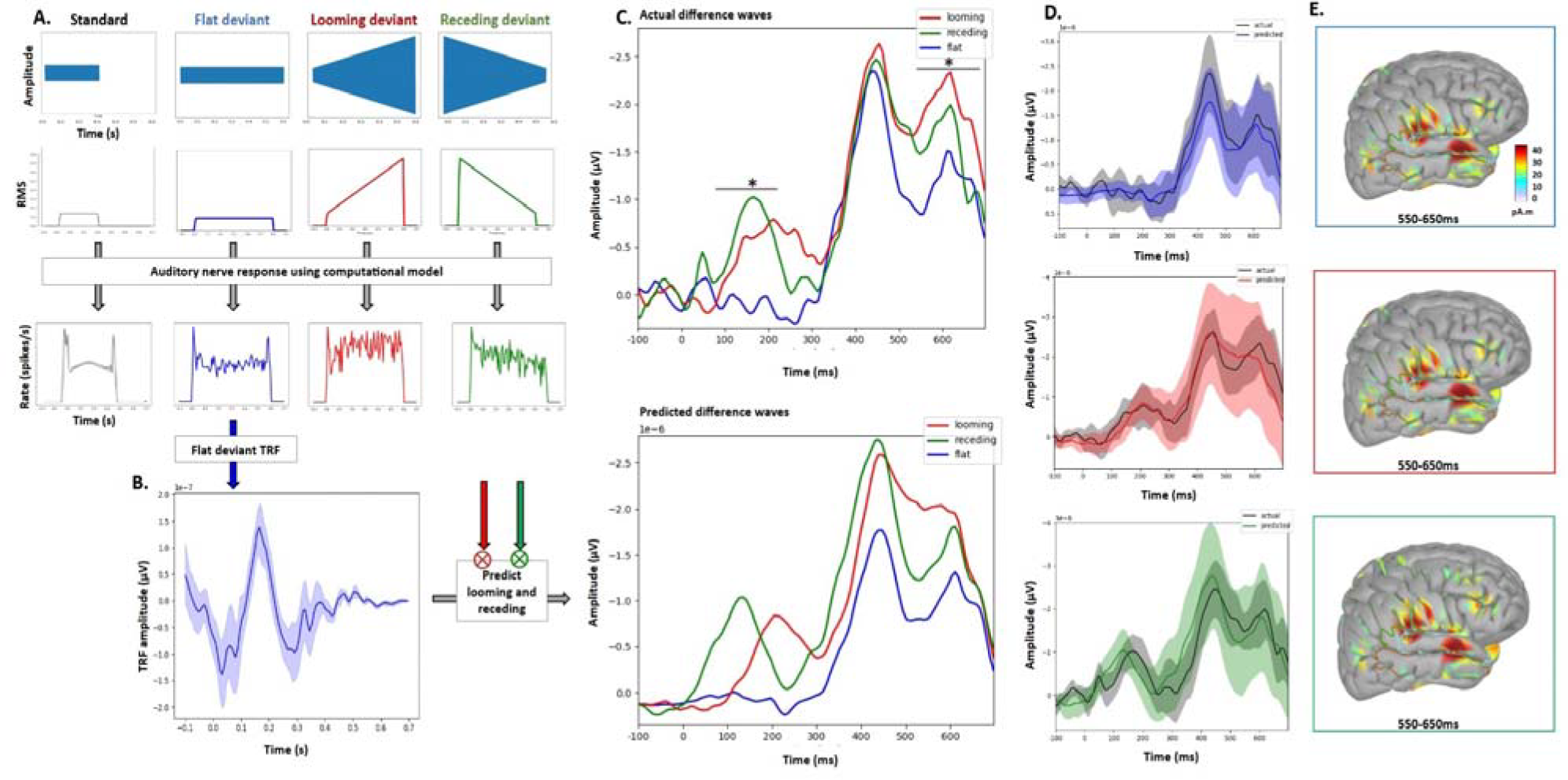
Cortical asymetries between looming and receding sounds are explained away by the auditory periphery. **A**. We collected scalp EEG mismatch negativity (MMN) data in response to deviant stimuli that had either dynamic (looming and receding) or constant (flat) level differences to the standard, in the same participants. Top: waveform and RMS intensity profiles for all four types of stimuli. Bottom: simulated auditory nerve response for the four types of stimuli, using the computational model of Zilany et al (2014). **B**. We then modeled the transfer function between the simulated auditory response and ERP difference wave for flat-intensity deviants (blue) using temporal response functions (TRFs), and used them to predict the effect of the same mechanism on other two stimuli (red: looming, green: receding). **C**. Predicted response to looming and receding sounds matched the observed difference waves almost perfectly (looming: 45%; receding: 33% explained variance). Top: Grand average of the observed difference waves (deviant minus standard) at the Fz sensor. Bottom: Predicted difference waves according to the flat TRF model. **D**. Cluster permutation test at the Fz sensor for observed vs predicted flat (top, blue), looming (middle, red), receding (bottom, green). No statistical difference was observed between the predicted responses and the observed responses for either type of stimuli. **E**. Sources localization in the right lateral cortical surface for observed flat (top), looming (middle) and receding (bottom) responsed. All three types of deviants generated a right anterior temporal source (superior and middle temporal gyri) and a right posterior temporal source (auditory cortices and planum temporale), again with no statistical difference between stimuli. Taken together, these results are evidence that MMN responses to looming and receding sounds result from the same cortical mechanism that generate MMN responses to unremarkable constant-level deviants.

At the scalp level, difference waves between deviant and standard sounds showed a striking succession of “MMN-like” negative deflections, which extended approximatively 170ms, 450ms and 600ms after stimulus onset (Figure 1.C). The first peak, which corresponded to a fronto-central midline distribution with maximum negativity at Fz (Figure 1.suppl material), was compatible with a MMN due to the initial difference between stimuli. Compared to *receding, looming* elicited a later MMN peak latency (206.5 vs 175.5ms, p=0.022) with a wider area under curve (AUC: 0.116 vs 0.053, p=0.002)(Table 1.supple material). Expectedly, there was no such peak for flat deviants, which did not differ from standards in that time range. The second peak, elicited between 400 and 500ms - thus about 150ms after the end of standard sounds, was compatible with a response to the end of standard sounds (Figure 2.suppl material). It occurred in a similar manner in all three types of deviants, with no statistical difference in either evoked responses characteristics (peak amplitudes, peak latencies and AUC) or surface amplitude maps. Finally, all three types of deviants elicited a late negative component between 550 and 650ms which, like the other peaks, had a fronto-central distribution with maximum negativity at Fz (Figure 1.suppl material). This late component also largely differentiated *looming and receding* sounds from flat sounds both in terms of peak amplitude, which was higher for *looming* (−1.96μV) than flat (−1.43μV, p=0.01); peak latency, which was earlier for *receding* (599ms) and *looming* (602ms) than flat (618ms, p=0.03); and AUC, wider for *looming* (0.263) than *receding* (0.197, p=0.05) and flat (0.181, p=0.023)(Figure 1.C and table 1.suppl material). In sum, at the scalp level, looming sounds displayed a clear pattern of MMN amplification, with wider responses in the range 150-250ms and earlier and more intense responses in the range 550-650ms. This pattern of result was consistent with other examples of looming amplification both in behavior (Neuhoff, 1998(2)), brain imaging (Seifritz et al. 2002(11)) and electrophysiology (Shestopalova et al 2018(13))).

To tease apart bottom-up auditory components from more specific top-down contributions in the response to looming and receding sounds, we then used generative EEG methods (temporal response functions, TRFs) to measure the extent to which the latter can be predicted from the former. First, we used a computational model of the auditory periphery (Zilany, Bruce & Carney, 2014(18)) to simulate the nonlinearities (i.e. loudness compression, temporal integration, onset and offset amplification) observed at the level of the auditory nerve, and how these affect the root-mean-square (RMS) profile of flat, looming and receding stimuli (Figure 1.A). We then used TRFs to model each participant’s MMN response to the output of this auditory model for flat-intensity deviants. TRFs provide an approximation of the mapping between incoming stimuli (here, the difference wave between the peripheral encoding of flat deviants minus standards) and the output EEG using a simple linear, time-invariant model represented by an impulse response (Crosse et al., 2016(19)). Here, we trained a separate TRF for each individual participant that predicts the participant’s MMN response to flat deviants (figure 1.B). The predicted response closely matched the observed response (80% explained variance; Figure 1.D), and thus provided an accurate model of the cortical response to a simple change of stimulus duration, at constant level. Finally, we used that flat-deviant model to simulate the extent to which bottom-up auditory mechanisms could explain the response to time-varying *looming* and *receding* stimuli (Figure 3.suppl material). To do so, for each participant, we convoluted the TRF trained on their flat response with the non-linear peripheral encoding of the two other types of deviants (*looming* and *receding*). Strikingly, both predicted responses almost perfectly matched the observed response (Figure 1.D): in particular, predicted *looming* responses exhibited the same pattern than seen in actual responses, i.e., a later/wider peak in the 150-250ms range, and an earlier/larger peak in the 550-650ms range. We tested for statistical differences between the predicted and observed responses across participants using cluster-based permutation test, and neither was significant. The flat model explained a remarkable 45% variance of the *looming* response, and 33% of the *receding* response.

Finally, we extracted cortical current source densities in the 150-250ms and 550-650ms time windows, and checked for significant differences between deviants. All three types of deviants generated a right anterior temporal source (superior and middle temporal gyri), a right posterior temporal source (auditory cortices and planum temporale) and to a lesser extend a right prefrontal cortex source (Figure 1.E). Finally, we also highlighted a left inferior temporal gyrus source (Figure 4. suppl material). None of them differed statistically, suggesting that they were generated by the same mechanisms (figure 5. suppl material).

In short, our results show that the large differences between cortical responses to *looming* and *receding* sounds exhibited at the scalp-level are in fact almost entirely explained away by non-linear encoding at the level of the auditory periphery, and result from the same cortical mechanism that generate MMN responses to unremarkable constant-level deviants. In other words, there is nothing cortex specific in the processing of *looming* sounds up to the level of the MMN response: all differences observed in MMN are the sole consequence of their particular physical morphology getting amplified and integrated by peripheral auditory mechanisms.

The sharp contrast seen here between the visually-salient scalp-level effects and their unassuming explanation by early peripheral differences should provide a sobering reminder that not all cortical effects, even in relatively late time-windows such as seen here, proceed from top-down modulations by high-level decision variables, such as a stimulus’ supposed physical, affective or social relevance. Rather, a lot of such differential amplification observed at the level of the cortex is in fact performed early and efficiently by feed-forward peripheral mechanisms that evolved precisely for the purpose of sparing subsequent networks with the necessity to implement such mechanisms.

## Methods

### Participants

N=18 volunteer participants (9 females, Median age = 25 years, Standard deviation SD=4.9, all right-handed) took part in the experiment. All participants reported no neurologic or psychiatric diseases and normal hearing. The experimental protocol was approved by Institut Européen d’Administration des Affaires (INSEAD)’s Institutional Review Board (protocol ID: 2021 – 51), and all participants gave written informed consent before the start of the study. They were financially compensated for their participation (25 euros/participant).

### Stimuli

1000Hz pure tones of different time-varying intensities were generated with custom Python software. Standard sounds had a duration of 300ms, with constant (root-mean-square) RMS intensity. All three types of deviants (flat, *looming* and *receding*) had a duration of 600ms (Figure 1.A). Flat deviants had the same constant RMS-intensity as standards. Looming deviants started at the same intensity as flat and standards, but their RMS-intensity increased linearly by 15dB over the duration of 600ms. Receding deviants started at the maximum intensity reached by looming deviants, and their RMS-intensity decreased linearly by 15dB over the duration of 600ms.

### Procedure

We used an *oddball paradigm* with standard (i.e., frequent and repetitive sounds) mixed with deviant (i.e., rare and unpredictable) sounds, the deviants being presented in a random order. The interval between the end of the sound and the beginning of the next one was set at 600ms. Standards represented 80% of all sounds, all deviants represented 20% (i.e., 6.7% of each), corresponding to 1601 standards and 133 deviants of each type. Three blocks of auditory stimuli were delivered for each participant, each block including 534 standards and 133 deviant sounds of all three types, with an inter-block interval duration of 5 minutes. Subjects were seated in a comfortable chair, in a quiet testing room and started to listen to the experiment. Sounds were presented binaurally over headphones, delivered by Python software. The participants were naive with respect to the hypotheses under test. Participants were asked not to pay particular attention to sounds, and to fix their attention on a relaxing mute video.

### EEG recording and data processing

EEG signals were recorded using a 64-channel EEG (actiCHamp, Brain Products GmbH, Germany). Data were continuously recorded with a high-pass filter at 1 Hz and a 1000 Hz sampling rate. EEG sensors were placed according to the 10-10 system (Seeck et al 2017(20)). Cz was set as the reference electrode. Sound onset triggers were sent to the EEG acquisition computer by a Cedrus StimTracker (Cedrus Corporation, San Pedro, CA). Preprocessing and EEG analyses were performed with EEGlab/Matlab R2022b (Delorme and Makeig 2004(19)) and replicated with Python the minimum norm estimate MNE (Gramfort et al, 2013(20). We first applied a band-pass filter between 0.1 and 30 Hz and a notch filter of 50 Hz (Parks-McClellan filter). We removed eye blinks and muscles artifacts using independent component analysis (ICA), after a 1Hz high-pass filtering (Delorme and Makeig, 2004(19)). We then interpolated the individual channels with artifacts using a 3-dimensional spine algorithm. EEG data were then re-referenced to the average reference and we segmented the EEG continuous data into epochs of 900ms, ranging from - 100 to 800ms relative to sound onset. We applied a baseline correction for each trial before stimuli onset (−100 to 0ms). All artifactual epochs, with voltage changes exceeding ±50 mV, were rejected from the analysis. Two subjects with more than 32% of epochs rejected were excluded from analysis, leaving 16 participants in the final analysis. ERPs of each participant were obtained by averaging separately each deviant and the standard stimuli using ERPlab software(Lopez-Calderon and Luck 2014(21)). Grand-averages were performed by averaging epochs for each condition in all participants.

MMN responses were obtained from the difference waveform between deviants and standards, and compared between our three deviant conditions (looming, receding and flat). Peak latencies, peak amplitudes and area-under-curves (AUC) were automatically measured at the Fz sensor, using EEGlab software. For each ERP component, a mixed analysis of variances (ANOVA) with the within-subject factor stimulus type (the three types of deviants) was calculated. Post-hoc comparisons were made using independent sample t-test. Differences were considered significant when p value was < 0.05. JASP software (JASP Team (2022), version 0.16.3) was used for statistical analysis.

### Auditory periphery modeling

In order to model the cortical response to *looming, receding* and flat deviant sounds, we used a computational model to simulate the effect of inner-ear-cell and auditory nerve (AN) non-linearities on the RMS profile of the three types of sounds. The model, described in Zilany, Bruce & Carney (2014) and implemented as a web application at https://urhear.urmc.rochester.edu/webapps/home/session.html?app=UR_EAR_2022a is one of two Auditory Nerve models to choose from: Zilany et al. (2014) and Bruce et al. (2018)(22,23). We used the model developed by Zilany et al. (2014) with human parameters for sharpness tuning and middle ear filter model provided by Ibrahim & Bruce (2010)(24). The Quick Plot functionality, which implements a single-CF Auditory Nerve model, was used to generate average discharge rate plots at a single characteristic frequency (CF) of 1000Hz for each of the three deviants.

### TRF analysis

To simulate the extent to which auditory non-linearities could explain the differential response to time-varying looming and receding stimuli, we used generative EEG methods (temporal response functions, TRFs) to model single-participant MMN responses to flat-intensity deviants, and used them to predict the effect of the same mechanism on time varying looming and receding stimuli. TRFs are impulse response functions that describe the relationship between the input and the output of a linear, time-invariant system. TRFs operate under the assumption that for a stimulus *s* at time *t* there exists a linear convolution with *s(t)* that results in the output of the system at that time *r(t)*. For a system with N recording channels, we can represent the neural response at time *t* and a specific channel *n* as the sum over all time lags *τ* of the linear convolution of stimulus characteristic *s(t)* and the TRF for that particular channel *w(τ, n)*.

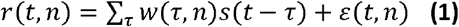

This linear convolution, referred to as the TRF, is estimated by minimizing the mean squared error between the actual and predicted responses using regularized ridge regression. In this study, we used the mTRFpy library (Crosse et al., 2016 (25)).

For each participant, we modelled the flat response using the following procedure: first, the AN response to flat sounds was subtracted by the AN response to standard sounds to get the stimulus difference (input). Second, all EEG epochs in which the flat deviant was presented were subtracted with the standard epoch preceding it to get the “flat-standard” EEG difference wave (output). We then estimated the input-output TRF by first, doing an exhaustive search for the best regularization parameter based on their cross-validated correlation between the predicted and measured response; and second, using the regularization parameter with greatest accuracy to build the final model. We obtained a single TRF for each participant. That TRF was then convoluted with the looming-standard and receding-standard stimuli to predict the neural response to the respective sounds. We ran temporal cluster permutation tests to test for any significant differences between the actual and predicted responses.

### Source localization

The estimation of cortical current source density was performed with Brainstorm (Tadel et al., 2011(24)). EEG electrodes positions were aligned to the standard Montreal Neurological Institute (MNI) template brain provided in Brainstorm. The mean head model was computed with the OpenMEEG Boundary Element Method for all participants (Gramfort et al 2010(26)). A noise covariance matrix was computed for each participant by taking the 100ms baseline period of each trial. For each subject, we computed one sensor-level average per condition (standard, *looming, receding* and flat). We then estimated sources for each average during the time window [-100; 800ms] using different methods of standardization: minimum norm estimate, dynamical Statistical Parametric Mapping (dSPM; Dale et al., 2000(27)) and standardized Low resolution brain Electromagnetic Tomography (sLORETA; Pascual-Marqui et al, 2002(28)). Source cortical maps were then compared with permutation paired t-test between the different types of sounds, in the different time-windows identified as significant in the grand average, and in ROIs recognized as of interest for MMN (Alho et al, 1995(29)).

## Data availability

The experimental datasets generated and analysed during the current study are available from the corresponding author on reasonable request.

## Abbreviation

AN: auditory nerve
AUC: area under curve
CF: characteristic frequency
dB: decibels
EEG: electroencephalography
ERP(s): event-related potential(s)
ICA: independent component analysis
MMN: mismatch negativity
ROIs: regions of Interest
RMS: root-mean-square
SD: standard deviation
TRFs: temporal response functions

