## supplemental file for "Cortical responses to looming sources are explained away by the auditory periphery"

**Supplementary materials.**

**Figure 1 supplementary material. The difference waves (“deviant minus standard”) at the Fz sensor, according to the *looming, receding* and flat conditions.** We observe three type of responses: a negative early component between 150 and 250ms, a second negative component between 400 and 500ms and a late negative component between 550 and 650 ms (upper part). Surface topographies of the difference waves, according to the type of deviant sound and to the early (around 200ms), second (around 500ms) and late (around 600ms) responses are presented in the lower part. At the bottom left, we observe a negative amplitude only for *looming* and *receding* sounds in the frontal areas, compatible with a MMN**.** At the bottom middle and right side**,** we observe a negative amplitude in the frontal area for all type of deviants.

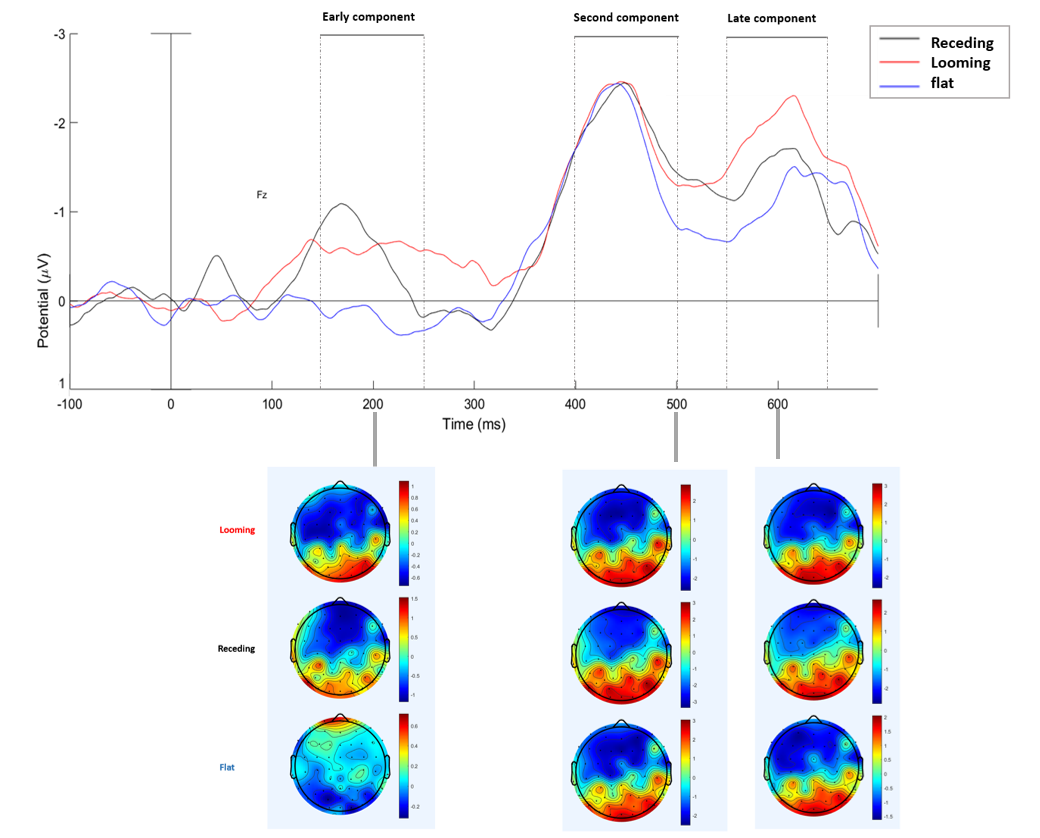

**Figure 2 supplementary material. Event related potential (ERP) grand-average for standard and deviant sounds at the level of Fz sensor. We highlighted the different evoked potentials (lower part) according to the type of stimuli onset (upper part).** We observe a negative component 100ms after stimulus onset for all type of sounds, compatible with a N100. We also observe a component at 150-250ms, with a visual waves mismatch regarding *receding* vs standard and *looming* vs standard sounds. There is no difference between flat and standard, because of the auditory characteristics of these sounds at this time. Around 400ms, there is a second ERP for the three types of deviants, compatible with a response to the “end of standard sounds”. We also can observe a late ERP around 600ms, and finally a last response at 750ms, namely 150ms after the end of the deviant sounds compatible with a response to the “end of deviant sounds”.

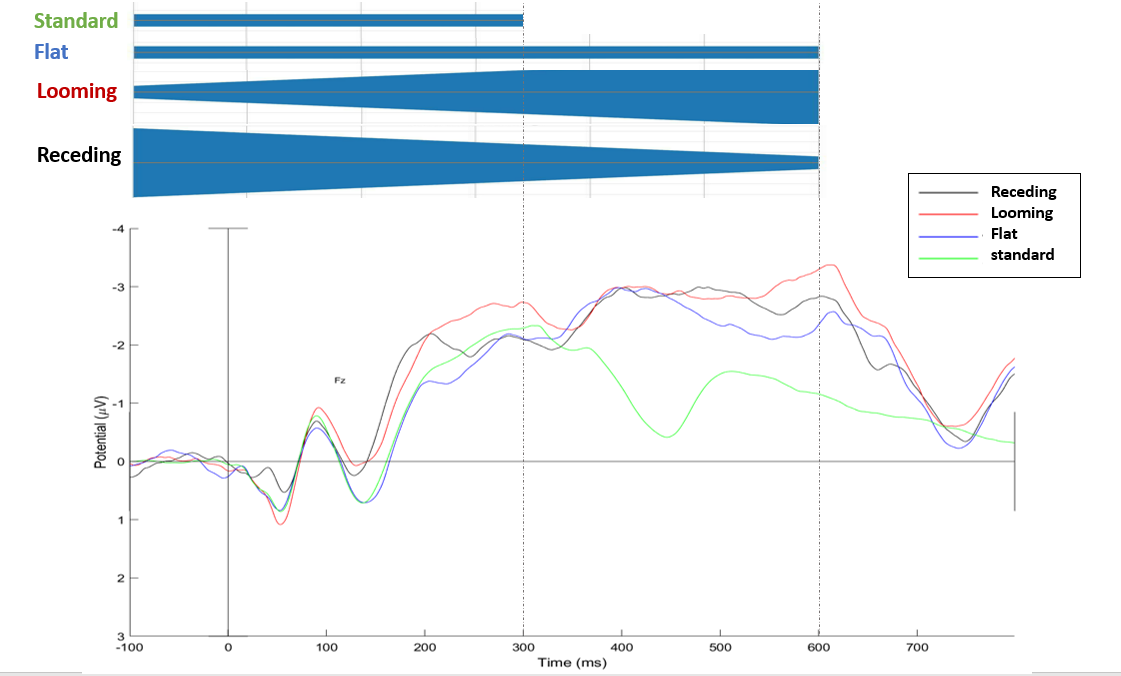

**Figure 3 supplementary material. TRF analysis.** For each participant, we modelled the flat response using the following procedure: first, the auditory nerve (AN) response to flat sounds was subtracted by the AN response to standard sounds to get the stimulus difference (input). Second, all EEG epochs in which the flat deviant was presented were subtracted with the standard epoch preceding it to get the “flat-standard” EEG difference wave (output). We then estimated the input-output TRF by first, doing an exhaustive search for the best regularization parameter based on their cross-validated correlation between the predicted and measured response; and second, using the regularization parameter with greatest accuracy to build the final model. We obtained a single TRF for each participant. This TRF was then convoluted with the *looming*-standard and *receding*-standard stimuli to predict the ERPs to these respective stimuli.

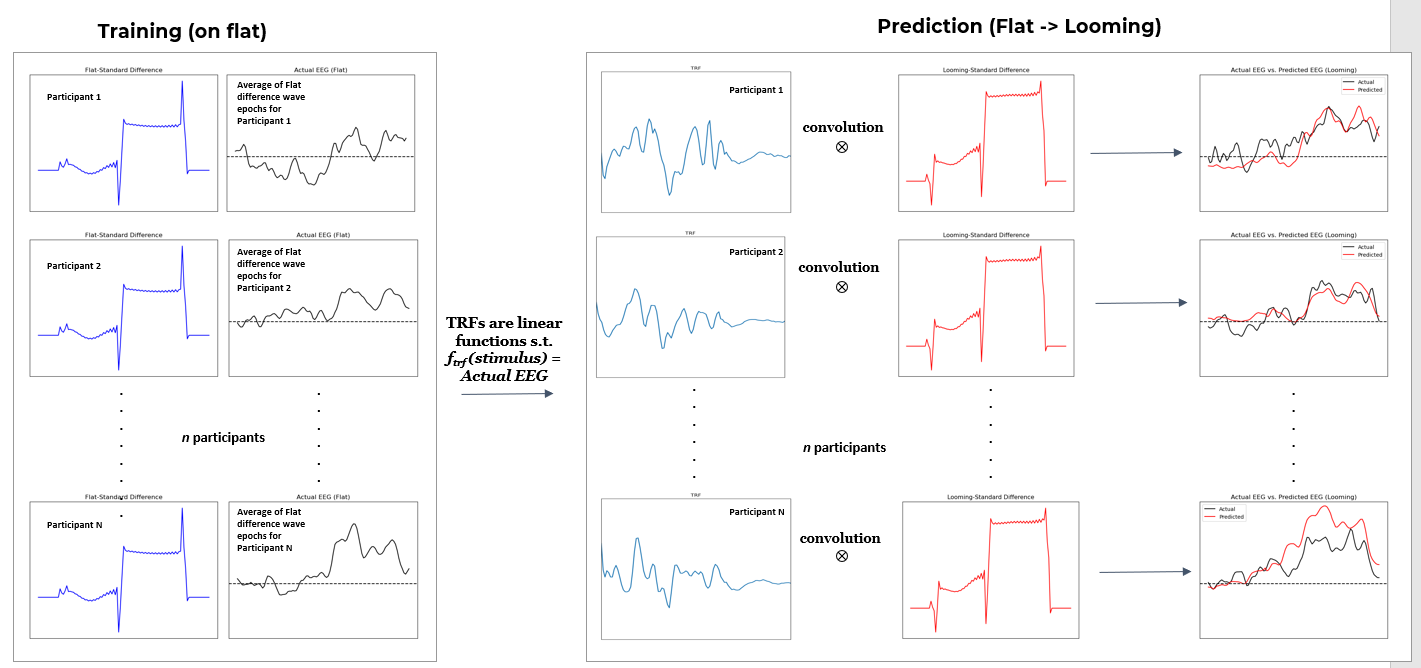

**Figure 4 supplementary material. Source localizations for standard, *looming, receding* and flat sounds.** The upper part shows the raw ERPs all 64 channels superimposed, for each type of sounds in the [-100;800ms] time window. The middle part of the figure shows the source localizations performed with the MMN (data option amplitude: 45%) at 200ms (early component) and the lower part the sources localizations at 600ms (late component). The different regions of interest (ROIs) identified by source localizations included in the statistical comparison permutation t-test are coloured: blue for the right supero-temporal; brown for the right middle temporal; green for the right supramarginal.

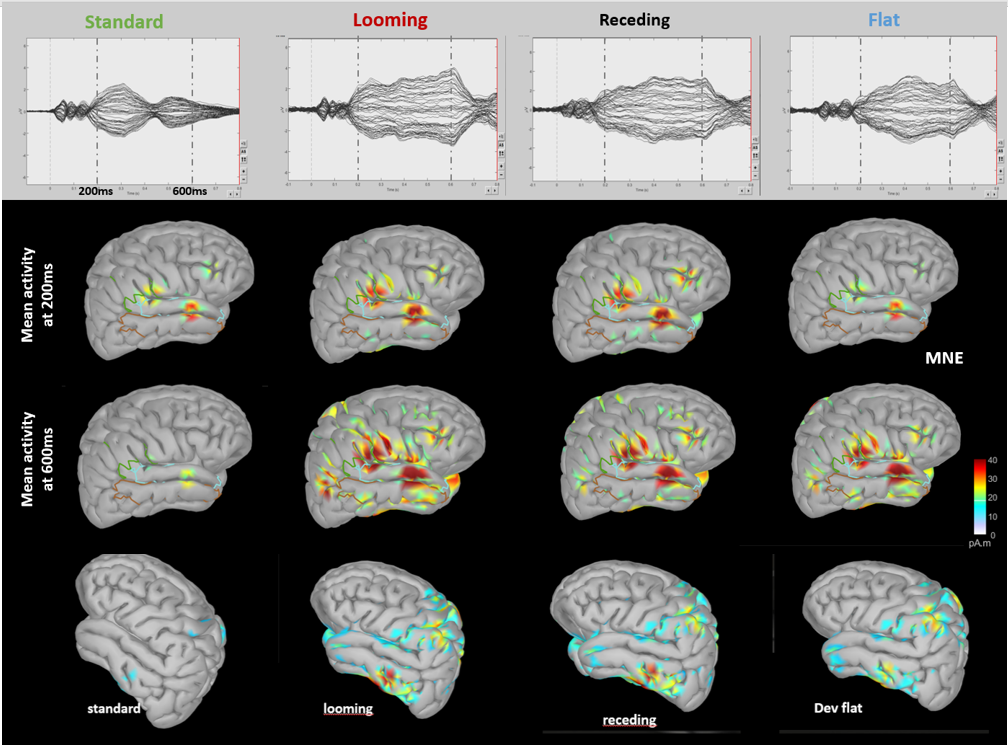

**Figure 5 supplementary material. Comparison of source localizations in the regions of interest (ROIs).** The left panel shows the marked ROIs in the looming condition in the two [150-250ms] and [550-650ms] time-windows of interest; The middle and right panels show the statistical analysis of sources comparison between conditions using permutation t-tests. The comparisons of sources revealed significant differences between standard vs *looming* and standard vs *receding* in the [350-650ms] time-window, but no significant difference between each type of deviant (i.e., *looming* vs *receding, recedi*ng vs flat and *looming* vs flat).

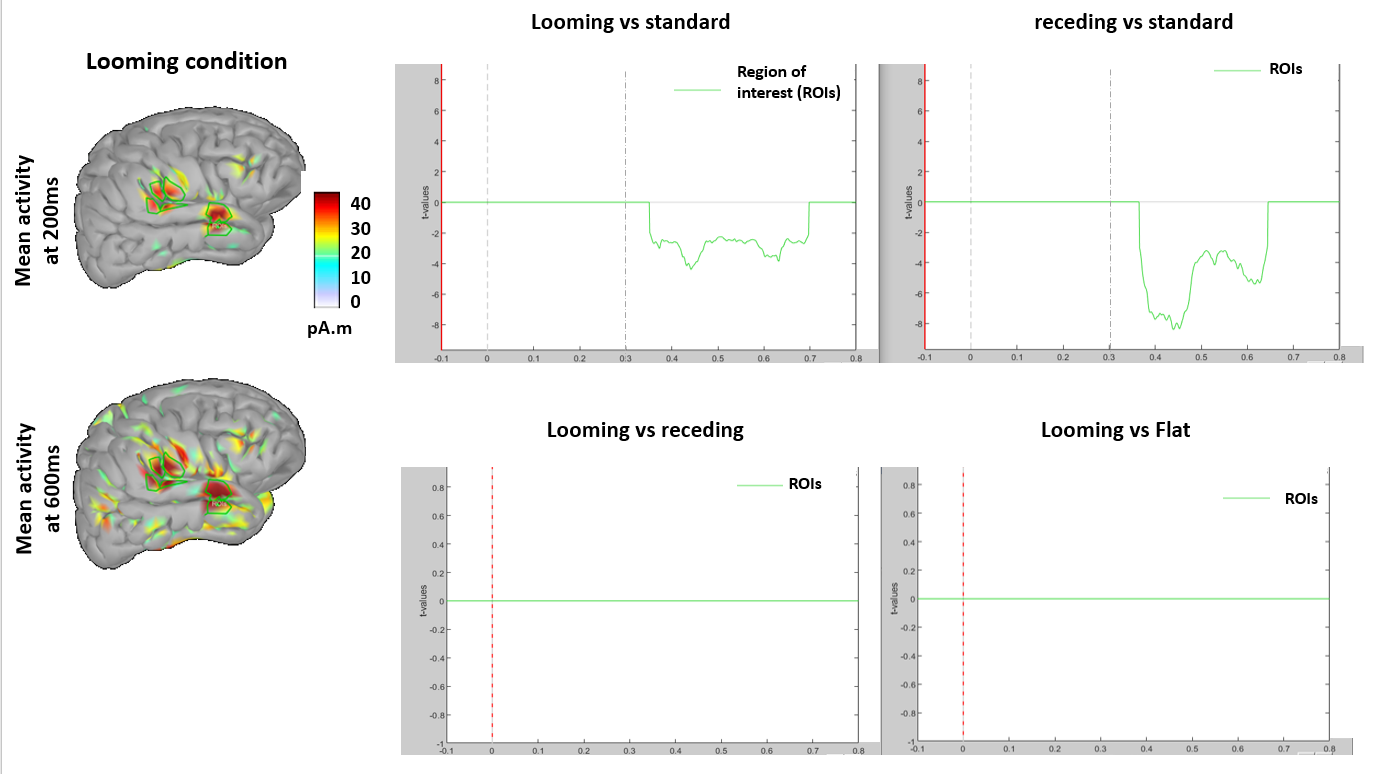

**Table 1 supplementary material. Comparison of peak amplitudes, peak latencies and area under curves (AUC) for the three “early” “second” and “late” components, according to the *looming, receding* and flat conditions.** Measures of peak amplitudes, peak latencies and AUC were performed at the level of Fz sensor. *indicate the significant difference with the p value, all other statistical comparison being not statistically significant. NA: not assessable; ms: milliseconds

| Conditions | *looming* | *receding* | Flat | *Looming* vs *receding* mean difference | *Looming* vs flat mean difference | *Receding* vs flat mean difference |
| --- | --- | --- | --- | --- | --- | --- |
| **Early component (150-250ms)** |  |  |  |  |  |  |
| Peak amplitude (µV, SD) | 1.24 (0.46) | 1.40 (0.98) | 0.21 (0.44) | 0.16  p=0.4 | NA | NA |
| Peak latency  (ms, SD) | 206.5 (29) | 175.5 (28) | NA | **31***  **p=0.022** | NA | NA |
| AUC (SD) | 0.116 (0.063) | 0.053  (0.063) | NA | **0.063***  **p=0.002** | NA | NA |
| **Second component (400-500ms)** |  |  |  |  |  |  |
| Peak amplitude (µV, SD) | 2.55 (0.98) | 2.50  (1.3) | 2.59  (1.4) | 0.05 | 0.04 | 0.09 |
| Peak latency (ms, SD) | 442.8  (18) | 445.4  (16) | 440.8  (17) | 2.6 | 2 | 4.6 |
| AUC (SD) | 0.315  (0.17) | 0.308  (0.21) | 0.285  (0.22) | 0.007 | 0.03 | 0.023 |
| **Late component (550-650ms)** |  |  |  |  |  |  |
| Peak amplitude(µV, SD) | 1.96 (1.2) | 1.61 (1.2) | 1.43 (1.33) | 0.35 | **0.53***  **p=0.01** | 0.18 |
| Peak latency (ms, SD) | 602 (24) | 599 (22.4) | 618 (17.6) | 0.03 | **16***  **p=0.03** | **19***  **p=0.03** |
| AUC (SD) | 0.263 (0.21) | 0.197 (0.19) | 0.181 (0.23) | **0.066***  **p=0.05** | **0.082***  **p=0.023** | 0.016 |
